# A Machine Learning-based Approach for Quantification of Protein Secondary Structures from Discrete Frequency Infrared Images

**DOI:** 10.1101/2025.01.08.632028

**Authors:** Harrison Edmonds, Sudipta S. Mukherjee, Brooke Holcombe, Kevin Yeh, Rohit Bhargava, Ayanjeet Ghosh

## Abstract

Discrete frequency infrared (IR) imaging is an exciting experimental technique that has shown promise in various applications in biomedical science. This technique often involves acquiring IR absorptive images at specific frequencies of interest that enable pathologically relevant chemical contrast. However, certain applications, such as tracking the spatial variations in protein secondary structure of tissue specimens, necessary for the characterization of neurodegenerative diseases, require deeper analysis of spectral data. In such cases, the conventional analytical approach involves band fitting the hyperspectral data to extract the relative populations of different structures through their fitted areas under the curve (AUCs). While Gaussian spectral fitting for one spectrum is viable, expanding that to an image with millions of pixels, as often applicable for tissue specimens, becomes a computationally expensive process. Alternatives like Principal Component Analysis (PCA) are less structurally interpretable and incompatible with sparsely sampled data. Furthermore, this detracts from the key advantages of discrete frequency imaging by necessitating acquisition of a more finely sampled spectral data that is optimal for curve fitting, resulting in significantly longer data acquisition times, larger datasets and additional computational overhead. In this work we demonstrate that a simple two-step regressive neural network model can be utilized to mitigate these challenges and employ discrete frequency imaging for retrieving the results from band fitting without significant loss of fidelity. Our model reduces the data acquisition time nearly 6-fold by requiring only seven wavenumbers to accurately interpolate spectral information at a higher resolution, and subsequently using the upscaled spectra to accurate predict the component AUCs, which is more than 3000 times faster than spectral fitting. Our approach thus drastically cuts down the data acquisition and analysis time and predicts key differences in protein structure that can be vital towards broadening potential applications of discrete frequency imaging.

## Introduction

Infrared spectroscopy-based chemical Imaging modalities like Fourier Transform Infrared (FTIR) imaging, Discrete Frequency Infrared (DFIR), and Optical Photothermal Infrared (OPTIR) Imaging have seen a resurgence in recent years as label-free imaging modalities that can augment conventional histopathology^1, 2^. In particular, the ability to map the chemistry of biospecimens for diagnostic and fundamental insights into disease pathologies has been galvanized by the development of discrete frequency-based approaches, wherein only the desired spectral bands of interest can be acquired for chemical contrast. As opposed to staining with dyes that differentially bind to different tissue components, chemical contrast is generated from the differences in their IR spectra arising out of biochemical compositions. These differences can then be exploited with unsupervised clustering methods^3^ or supervised learning algorithms^4^ to generate label-free segmentation of tissue histology. While significant spectral differences exist between cellular (epithelial cells, malignant cells) and extracellular components (stroma, secretions), subtle differences in Amide I line shapes arising from protein secondary structural variations can also be clinically important and are increasingly recognized for their pathophysiological significance. The characterization of protein bands is of relevance for disease pathologies involving amyloid aggregates, collectively termed amyloidosis.

Amyloidosis contributes significantly to neurodegenerative diseases like Alzheimer’s, Parkinson’s, Huntington’s, and Chronic Traumatic Encephalopathy (CTE) through the accumulation of misfolded protein aggregates. Each of these diseases involve aggregates of a specific proteins that misfold to form amyloid fibrils. In Alzheimer’s disease, Aβ plaques and neurofibrillary tangles disrupt synapses and intracellular transport, leading to cognitive decline^5–7^. In Parkinson’s, alpha-synuclein aggregates form Lewy bodies, impairing mitochondrial function and killing dopaminergic neurons, which causes motor dysfunction^8, 9^. Huntington’s involves mutant huntingtin protein aggregates that interfere with transcription and metabolism, leading to progressive neuronal loss in the striatum^10, 11^. In chronic traumatic encephalopathy (CTE), repeated trauma promotes tau aggregation and neuroinflammation, damaging neurons and causing cognitive and behavioral symptoms^12, 13^. Across these diseases, and more^14, 15^, misfolded proteins disrupt cellular function, trigger neuroinflammation, and propagate in a prion-like manner^16, 17^, under the term protein aggregation^18, 19^.

Infrared spectroscopy-based imaging approaches have been applied to study these pathologically relevant protein aggregates ^7, 20–24^, and these studies underscore the importance of quantifying protein secondary structural distributions for gaining key insights into the molecular biology of the disease. Traditionally, quantification of protein secondary structure from Amide I IR spectra is achieved by deconvolving the spectra into a predetermined number of Gaussian peaks (usually three to four), where the relative peak intensities, linewidths, and central frequencies are extracted with a constrained non-linear least square curve-fitting approach. The area under the curve (AUC) of the peaks can subsequently be interpreted as proportional to the populations of the constituent species, assuming similar transition dipole moment strengths^25^. Although the Gaussian-fitting method is relatively straightforward for a few spectra, it can become prohibitively computationally expensive when scaled up for whole-slide images (∼10-100 million spectra per patient) and is susceptible to convergence issues for noisy data^26^. As a result, only a tiny fraction of IR imaging studies have used spectral deconvolution via band fitting to get structural insights into proteins^27, 28^. Alternatives to deconvolution approaches such as Principal Component Analysis (PCA) are used more commonly^29–32^; however, the results from such analyses cannot be readily interpreted from a structural perspective^25^ and are incompatible with sparsely sampled IR/photo-thermal data^33^ often employed to circumvent the data acquisition overhead for a large specimen^34^. Secondly, spectral band fitting negates the advantages of discrete frequency-based approaches because it necessitates hyperspectral data acquisition. Since the experimental time for data acquisition linearly scales with the number of bands, such an approach becomes unfeasible for generating a statistically significant data set containing a large cohort of patients. Neural networks, particularly convolutional neural networks (CNNs), have shown great promise in the classification and segmentation of complex images. In the context of infrared spectroscopic imaging^1, 35–38^, CNNs can be trained to accurately identify and delineate pathological lesions for a variety of tissue samples, by leveraging both visual and near-infrared spectroscopic data^39^. Neural networks (NN) can also optimize the process of data collection. By using image up-scaling models, NNs can improve the spatial and spectral resolution of the collected images^40^, allowing high-resolution data to be obtained from lower-resolution images, thereby reducing the need for extensive and time-consuming data collection. This is particularly beneficial in medical imaging, where high-resolution images are crucial for accurate diagnostics but can be difficult and expensive to obtain^41–43^. However, efforts to upscale chemical images have largely focused on the spatial dimension^41^, while the importance of the spectral dimension has somewhat gone unnoticed. The data acquisition challenges detailed above can be significantly mitigated if spectral upscaling can be implemented in addition to spatial resolution enhancement, wherein only a handful of bands can be used to reconstruct finer sampled spectral data. This advancement allows for discrete frequency imaging to be applied in applications that typically require spectral deconvolution, eliminating the need to acquire a full spectrum for each pixel. Even for applications that traditionally rely on full spectra, this method demonstrates that they can be retrained to use inferred spectra from a few measured bands, streamlining the process and enhancing efficiency.

It is important to note in this context that several analysis methods besides band fitting can be employed to extract information about protein secondary structure from IR spectra. These include two-dimensional correlation spectroscopy, Fourier self-deconvolution and second derivative analysis. The latter involves calculating the second derivative of the IR spectrum, which enhances the separation of overlapping peaks and reveals subtle features that may not be apparent in the original spectrum and has been widely applied towards analysis of protein IR spectra^44–46^. Fourier self-deconvolution techniques aim to enhance the resolution of the IR spectrum by removing instrumental broadening and separating overlapping peaks by use of apodization in the Fourier transformed time domain signal. This allows for a more accurate determination of the number and positions of underlying peaks, which can then be assigned to different secondary structures^47, 48^. Two-dimensional correlation spectroscopy generates maps that reveal positive and negative correlations between different spectral components, which in turn reveals insights into the dynamic behavior of molecules and their interactions^49^. These approaches can also be augmented with other complementary approaches such as circular dichroism spectroscopy to provide a more nuanced and comprehensive picture of protein secondary structure^50^. However, all the above strategies rely on acquisition of a full spectrum. Our approach enables reconstruction of the entire spectrum from sparsely sampled bands, which can consequently be analyzed using any method as appropriate and thus can expand the applicability of these analytical approaches in spectroscopic imaging further. The second neural network reported in this study aims to reproduce the AUCs as typically determined from curve fitting; in principle other networks can be envisioned and trained to predict the parameters of different spectral analysis methods. We hope to address this in future work.

In this work, we implement a two-step NN regression-based approach to perform spectral reconstruction and extract the appropriate fit parameters from sparsely sampled discrete frequency IR images with an aim for overcoming the aforementioned limitations of spectroscopic imaging. Our approach utilizes two distinct models, the first of which takes a sparsely sampled seven-wavenumber spectral input of key spectroscopic frequencies selected in order to preserve the maximum amount of spectroscopic information and up-samples it to a forty-one-wavenumber (4 cm^-1^ spectral resolution) output, which is subsequently forwarded into an AUC prediction model. We first train NN models with simulated data with underlying distribution analogous to experimental data. Subsequently, we forward the model to various selections of experimental data. The advantage of our method over the traditional Gaussian fitting approach is two-fold. Firstly, the computational time needed to forward the pre-trained model is up to 3100 times faster than the conventional approach. Secondly, by only acquiring these seven select discrete frequency bands instead of a hyperspectral dataset, we cut the experimental scan time by approximately a factor of 6.

## Materials and Methods

### Tissue Samples

The study utilized 3 formalin-fixed paraffin-embedded (FFPE) diseased frontal lobe tissue samples from Alzheimer’s disease (AD) patients acquired from Advanced Tissue Services (Tucson, AZ), and 1 diseased temporal lobe tissue sample from BioChain Institute Inc. (Newark, CA). These postmortem tissue samples were deidentified and classified as non-human subjects research by the University of Alabama’s Office of Research Compliance. Before infrared (IR) imaging, the tissues underwent a 24-hour deparaffinization process in n-hexane and were stored under mild vacuum conditions. Adjacent tissue sections were used for immunohistochemistry (IHC) staining with anti-amyloid MOAB-2 antibody (Sigma Aldrich)^51, 52^ and silver staining^53^ to assist in identifying the location and morphological characteristics of amyloid-beta (Aβ) plaques.

### IR Imaging

A custom-built stage scanning infrared (IR) microscope using a quantum cascade laser system (Block Engineering; LaserTune) was utilized to capture IR images^54, 55^, This microscope offers tunable illumination ranges from approximately 1000 to 1800 cm^−1^. IR imaging was done in transflection mode on IR reflective low-emissivity slides (Kevley Technologies; MirrIR)^56^ from 1584 to 1730 cm^−1^, with a frequency resolution of 4 cm^-1^, with a 0.71 NA objective, and a pixel size of 2 μm.

### Network Construction

The neural network models were trained using simulated data containing 200,000 spectra of the Amide I band spectral region (1584 cm^-1^-1744 cm^-1^). The synthetic spectral dataset was split in half prior to training the two models: 100,000 were used to train the sparse data up-scaling model while 100,000 were used to train the AUC prediction model. The spectral up-scaling training dataset was augmented with 20,000 additional spectra from Alzheimer’s disease brain tissues. To sample the spectral heterogeneity of tissue infrared spectra, the training dataset was designed to encompass varying levels of noise, non-zero baselines arising from scattering effects, and variations of central peak positions and spectral widths. The spectral up-sampling model was trained using the sparsely sampled seven wavenumber input (1588 cm^-1^, 1612 cm^-1^, 1636 cm^-1^, 1660 cm^-1^, 1684 cm^-1^, 1708 cm^-1^, 1732 cm^-1^). It is important to note that the choice of the specific seven wavenumbers is not predicated on their association with specific secondary structures; rather, these wavenumbers constitute optimal spectral bands necessary for accurate reconstruction of a complete spectrum. This model contains three hidden layers with 16, 32, and 64 hidden neurons and an output layer which contains 41 neurons corresponding to a full Amide I bands spectral region with a spectral resolution of 4 cm^-1^. A schematic representation of this model is shown in Figure S5. This output is then used as input to the second model, which has two hidden layers with 16 and 9 neurons respectively, and an output layer with 3 neurons corresponding to the 3 AUC values. A schematic representation of this model is shown in Figure S4. All the neurons in our model were fully connected. We used a Scaled Exponential Linear Units (SELU) activation function for the hidden layer neurons in the AUC prediction model and a RELU activation function for the output layer neurons and hidden neurons within the spectral up-sampling model. The weights and biases of the networks were trained using the Adaptive Moment Estimation (ADAM) algorithm, and the AUC prediction model used LeCun normal initialization of weights within the network’s hidden layer to minimize the Mean Absolute Error (MAE) loss function. All training was performed with Python (3.8.17), and TensorFlow (2.13.0).

### Gaussian Fitting

Experimental and simulated spectra were linear baseline corrected and fitted to three Gaussian peaks roughly centered at 1632 cm^-1^, 1666 cm^-1^, and 1690 cm^-1^ with a range value of 5 cm^-1^ using the native Gaussian fitting option in MATLAB (nonlinear least square fit with Levenberg-Marquardt Algorithm) to extract the peak parameters for the three bands, fitting parameters detailed in Table S2.

## Results and Discussion

As summarized in Figure 1, we simulate 200,000 spectra as the sum of three intensity normalized Gaussian peaks with known AUC values. 20,000 additional spectra from Alzheimer’s disease brain tissue data were also added to the training dataset for the spectral up-scaling model. It has been shown previously that FFPE brain tissue Amide I spectra can be adequately deconvoluted as a sum of three bands^24, 46^; thus, the choice of three Gaussian bands for our model is suitable. Subsequently, 120,000 spectra were split randomly into training (80%) and testing (20%) datasets and used to train and evaluate each model respectively. The spectra were discretely sampled at 1588 cm^-1^, 1612 cm^-1^, 1636 cm^-1^, 1660 cm^-1^, 1684 cm^-1^, 1708 cm^-1^, and 1732 cm^-1^ to generate the input dataset for training the spectral up-scaling model with the densely sampled 41 wavenumber spectra as the target. Following the training of the upscaling model, the second set of 100,000 spectra were also split randomly into a training (80%) and testing (20%) and sparsely sampled at the seven wavenumbers previously mentioned and forwarded through the upscaling model (ANN1); the resulting output is then used to train an NN-regression model (ANN2) with the AUC values as the target. The combined pre-trained network is then forwarded on actual experimental data and the extracted AUC values are compared to those obtained from the conventional fitting approach.

**Figure 1:**
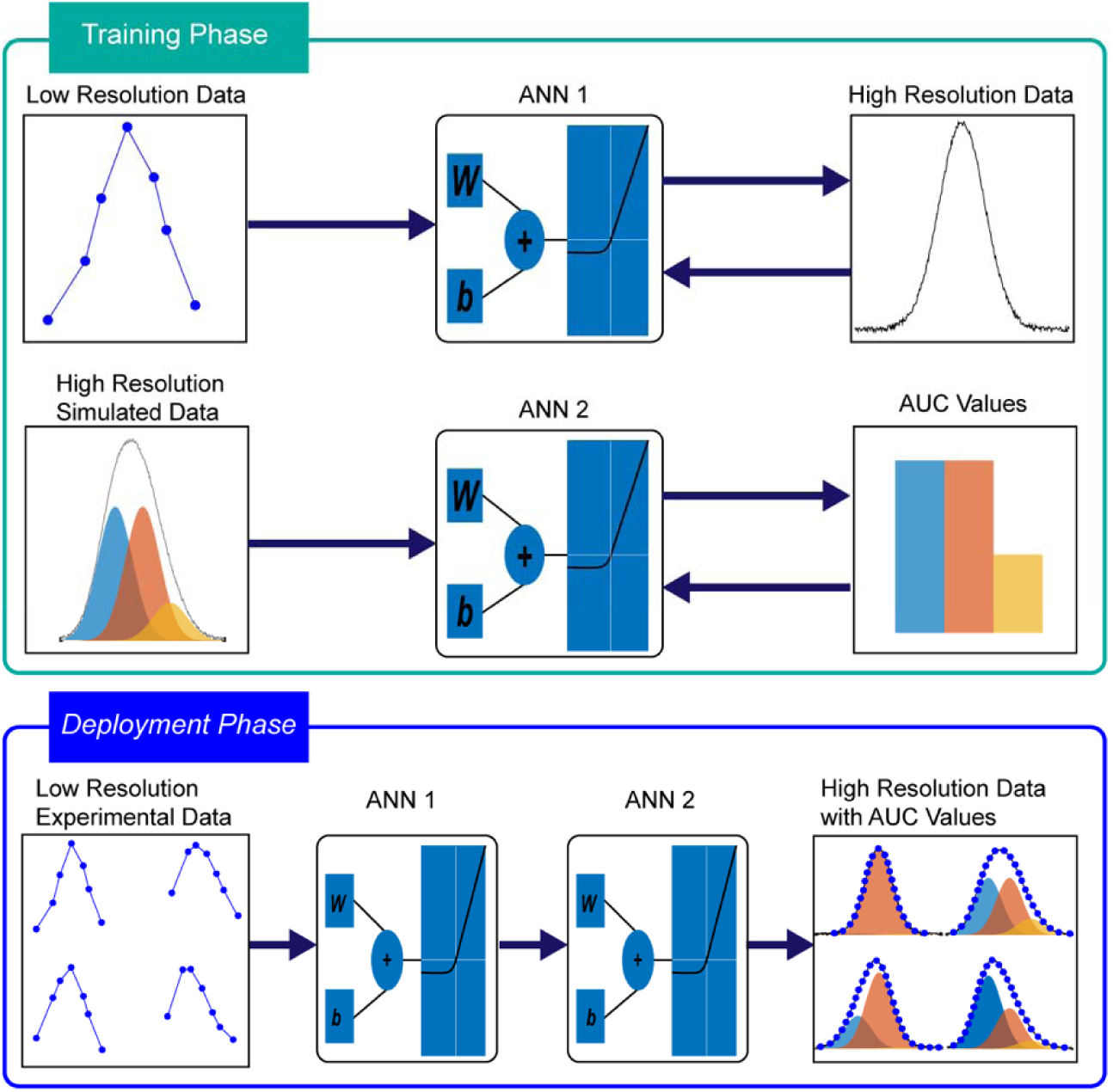
Schematic representation of the method presented in this paper. AUC: Area Under Curve (AUC1: Blue, AUC2: Orange, AUC3: Yellow), ANN 1: Artificial Neural Network responsible for up-scaling of spectroscopic data, ANN 2: Artificial Neural Network responsible for estimating the deconvolution of said upscaled spectroscopic data.

To account for absorbance variations between tissues originating from thickness and other artifacts, the input to the neural networks were mean intensity normalized. Figure 2 highlights the accuracy of the spectral upscaling model by displaying three representative examples of actual vs. model predicted spectra (2A) along with an example of images from amyloid plaques, a key morphological feature in Alzheimer’s disease^57–59^. An infrared spectral image from this plaque was collected at 1628 cm^-1^ and displayed below (2C). This example agrees well with the corresponding upscaling model output predicted for 1628 cm^-1^, resulting in a Structural Similarity Index Measure (SSIM) of 0.9969. The spectral upscaling model was also evaluated using the test dataset and was found to have a 0.015 MAE. To optimize the accuracy of our spectral upscaling model with the speed of experiment and analysis, we evaluated the dependence of MAE on the number of input bands to the AUC prediction model (Figure S1). This analysis indicated clear diminishing returns in terms of model performance when training with additional wavenumbers beyond 7. During this series of tests, the Amide I band was undersampled with 7 uniformly spaced measurements. The final model uses 7 wavenumbers that are not equally spaced but are rather chosen better to sample the characteristics of the 3 underlying bands. We verified that this specific wavenumber set slightly improves the model accuracy compared to using 7 equally spaced bands and the results are shown in Figure S1, where the red dot represents our modified selection of wavenumbers and the corresponding 7 wavenumber blue dot represents 7 equally spaced wavenumbers.

**Figure 2:**
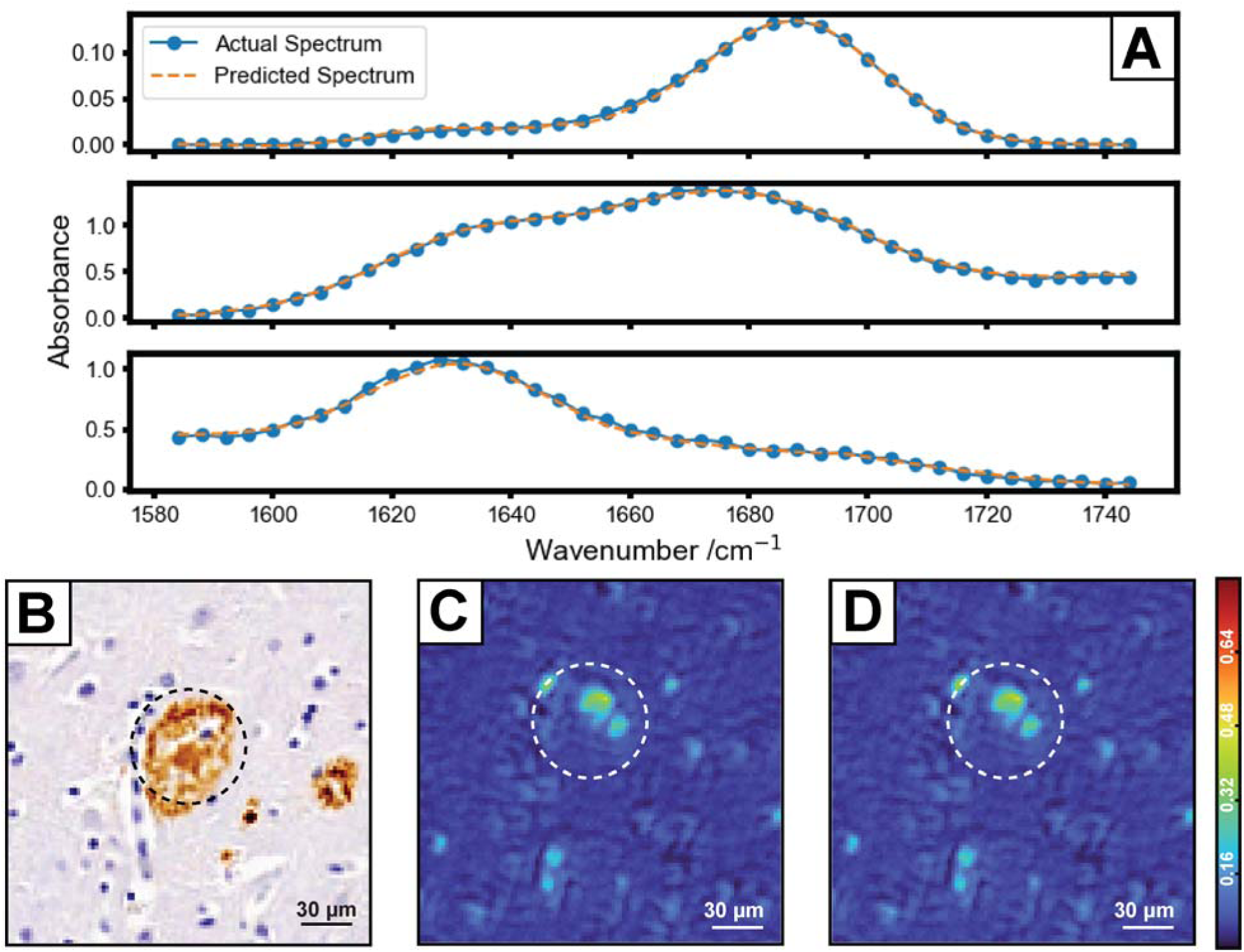
(A) Various example spectra from the test dataset and their corresponding predicted spectral output, (B) optical IHC stained Aβ plaque, (C) intensity image collected at 1628 cm^-1^ of the plaque corresponding plaque with the approximate position estimated by the white dashed circle, (D) the same intensity image predicted by model output.

Upon validating the fidelity of the predicted spectra, we applied the second model to the spectra generated from Model 1 to predict the AUCs of the constituent bands. Figure 3 displays the simulated dataset’s predicted AUC vs ground truth AUC and the corresponding R^2^ values and linear fits. The high R^2^ values and small intercepts (relative to the mean AUCs) validate our AUC prediction model’s performance. The MAE of our test set evaluated for all three AUC values was 5.16. Out of the three fitted peaks, the lowest R^2^ value is for AUC2 (0.880) because of the possibility of overlap from the two Gaussian peaks on either side. This is a well-known issue with deconvolving broad spectra and arises due to the extent of fundamental ambiguity in the deconvolution approach^60^. By leveraging this model, the time required for spectral acquisition is cut dramatically. The experimental time for discrete frequency imaging typically scales linearly with the number of bands, meaning measuring a sparse seven-wavenumber dataset compared to the typical forty-one wavenumbers required to investigate the slightly expanded Amide I band spectral region (1584-1748 cm^-1^) with 4 cm^-1^ spectral resolution leads to a roughly 6-fold decrease in experimental time For biological samples where it is essential to sample the interpatient heterogeneity for statistically significant results, such reduction of experimental and computational times is invaluable.

**Figure 3:**
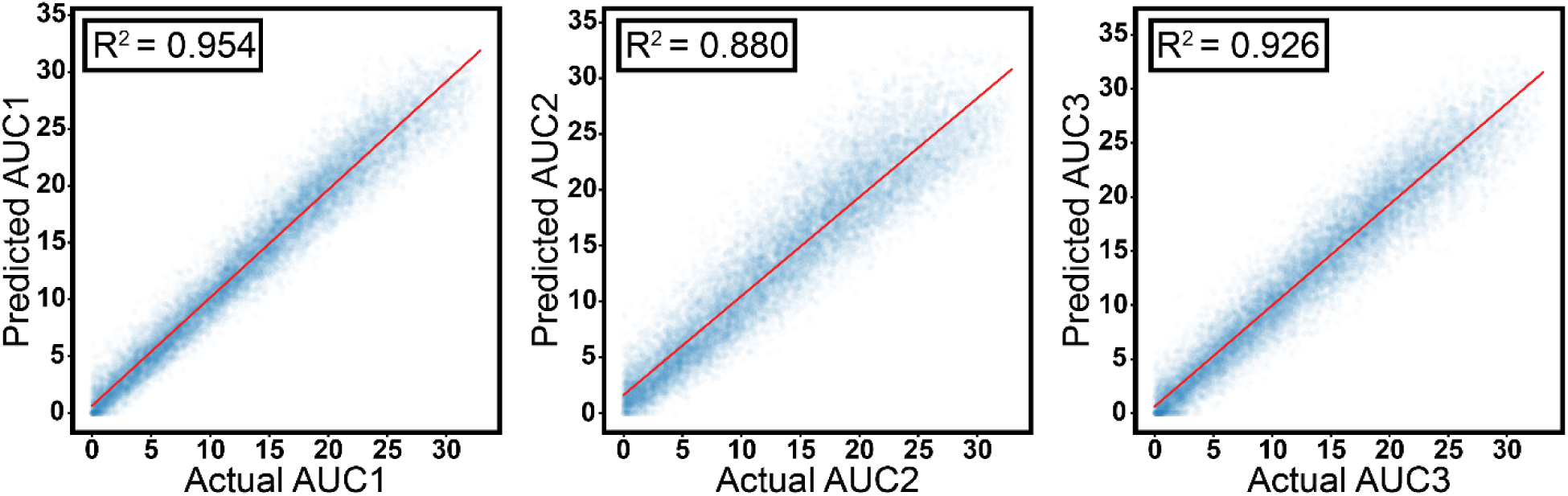
Scatter plots of Actual vs Predicted AUC values corresponding to the test dataset split.

After the training phase, the model’s performance was evaluated on experimental hyperspectral images from Alzheimer’s disease brain tissue specimens. Neuropathological diseases such as Alzheimer’s, Parkinson’s, etc., involve the aggregation of specific proteins into extra and intracellular deposits, which are believed to play important roles in the development and progression of the disease. However, protein structures in the brain are yet to be fully understood, and how these aggregates truly influence disease pathology at a molecular level is unclear. In particular, how the secondary structure of these aggregates evolves with disease pathology and if/how that parallels in-vitro studies remains a topic of intensive investigation. Accurately mapping protein secondary structure requires spectral deconvolution, making our model ideal for such applications. To compare the model output with the current gold standard in spectral deconvolution, namely band fitting, the infrared Amide I band spectral contribution collected from measuring these plaques were deconvolved into three Gaussian peaks with both the model and Gaussian fitting algorithm. These three Gaussian peaks are centered at approximately 1628 cm^-1^, 1660 cm^-1^, and 1690 cm^-1^ the integrated Area Under the Curve (AUCs) of these three peaks will be referred to as AUC1, AUC2, and AUC3 respectively in this article. Integrating each peak allows for a comparison of the relative strength of each from one pixel to the next, in the context of protein spectroscopy the relative abundance of AUC1 is of particular importance, since the 1628 cm^-1^ peak correlates to relative parallel β-sheet concentration^61^ which can be predictive of many key neurodegenerative morphological features like Aβ plaques. Figure 4A&D shows the optical images of the IHC stained parallel section for the two plaques, while Figure 4B&E and Figure 4C&F show the spectral fitting and network outputs, respectively. We can observe a clear correspondence between the fitted AUC1 images (4B&E), and network-measured AUC1 images (4C&F). An important distinction between the fitted AUC image and the Model AUC image is the difference in baseline (non-beta sheet values), where the Gaussian fitting presents significantly more background noise than the Model output. This issue is well-documented in the context of Gaussian fitting where the fitting process can amplify noise and create artificial peaks in the background, leading to misinterpretations of the spectral data. Gaussian fitting often struggles with distinguishing between actual signal and background noise, particularly in complex biological samples where spectral overlap and variations are common^62^. In the characterization of amyloid plaques, it is generally mitigated by concentrating on the region containing the plaque. However, increased image noise complicates the identification of the desired morphological feature.

**Figure 4:**
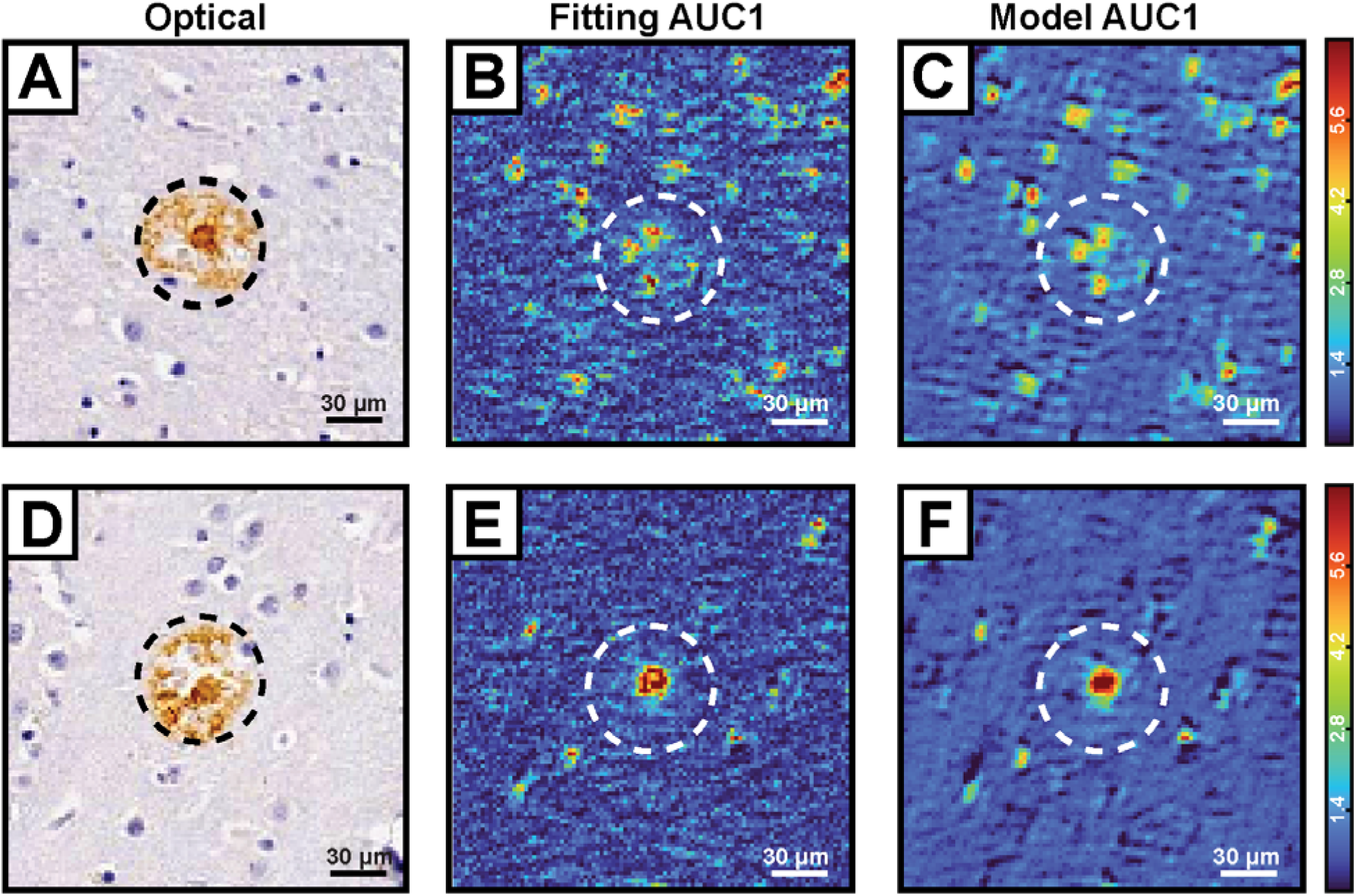
(A,D) Optical IHC stained images of Aβ plaques with the plaque identified by the black dashed circle, (B, E) deconvoluted AUC1 images from collected hyperspectral data and (C, F) reconstructed AUC1 images from network predictions with the approximate position of the plaque estimated by the white dashed circle.

The high versatility of our model also allows it to be utilized for much larger specimens, where the acquisition of a hyperspectral dataset can become prohibitively time-consuming. Furthermore, the size of such datasets also poses a challenge for analysis. Spectral fitting for such data would become a computationally expensive process that can significantly strain or even overwhelm available computational resources. Figure 5A displays AUC1 images reconstructed from the neural network model output of an entire frontal lobe tissue section. Figure 5B and Figure 5C display the Gaussian fitted and Model predicted AUC1 values respectively within a smaller hypothetical area of interest indicated by the dashed white line in Figure 5A, which clearly demonstrates the fidelity of our model when compared to spectral fitting. Figures S2-S4 present additional examples of AUC1 images reconstructed from the neural network model output.

**Figure 5:**
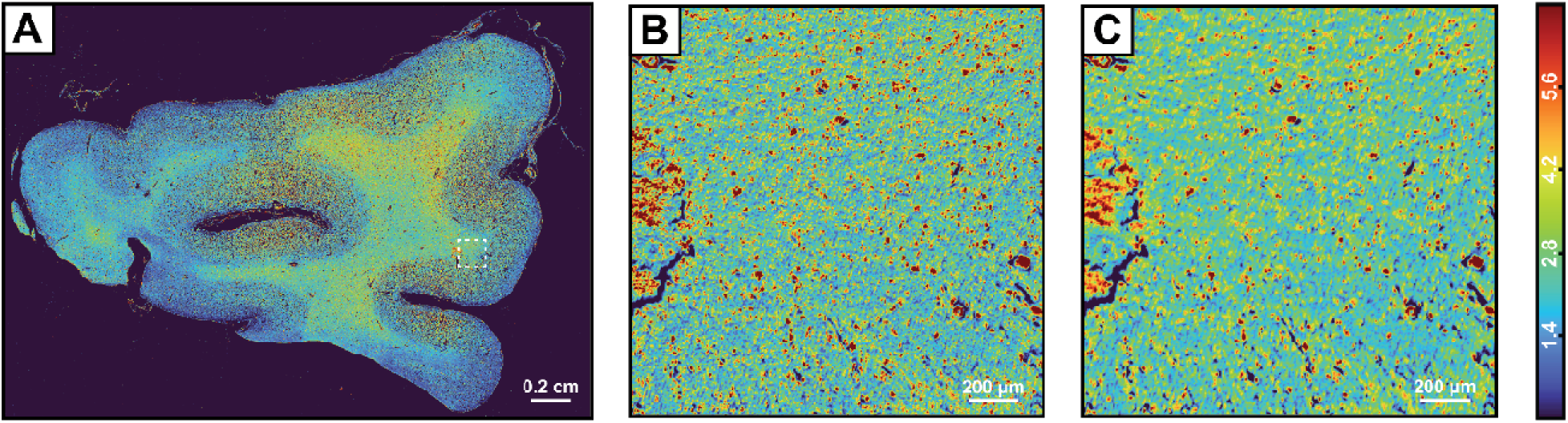
(A) Network predicted AUC1 image from collected hyperspectral data of a frontal lobe Alzheimer’s tissue section, (C) a smaller hypothetical region of interest of the same image denoted by a dashed white box, (B) the same cropped region of interest where the spectra were fitted using Gaussian fitting.

A single band at a 2 μm/pixel resolution for this specimen requires ∼67 minutes to acquire and results in an image with approximately 154 million pixels. Acquisition of a hyperspectral dataset optimal for subsequent deconvolution would thus require nearly 2 days and lead to approximately 6.3 billion pixel measurements. Subsequent spectral analysis of a dataset of this size is also expected to be significantly time and resource consuming. Our model allows for the reproduction of the spectral features and subsequent deconvolution with high fidelity and approximately 6-fold faster data acquisition without additional computational overhead.

Finally, the model prediction performance on simulated spectra with varying levels of added random noise was compared to the accuracy of the Gaussian fitting method outlined previously. The results of this analysis are presented in Figure 6 (tabulated in Table S1) and indicate that the model consistently outperforms Gaussian fitting. Figure 6 shows the relative MAE of the two deconvolution methods vs. the signal-to-noise ratio for various sets of simulated spectra; this value was assessed in the simulated datasets by calculating the standard deviation of the signal at the tail of the Amide I region (1742 – 1748 cm^-1^) where there is no expected signal. The noticeable difference in the slopes of the corresponding models in Figure 6 suggests that Gaussian fitting is much more susceptible to an increase in noise than our model. This agrees well with current literature, which suggests that Gaussian fitting has difficulty fitting to lower SNR data^62^.

**Figure 6:**
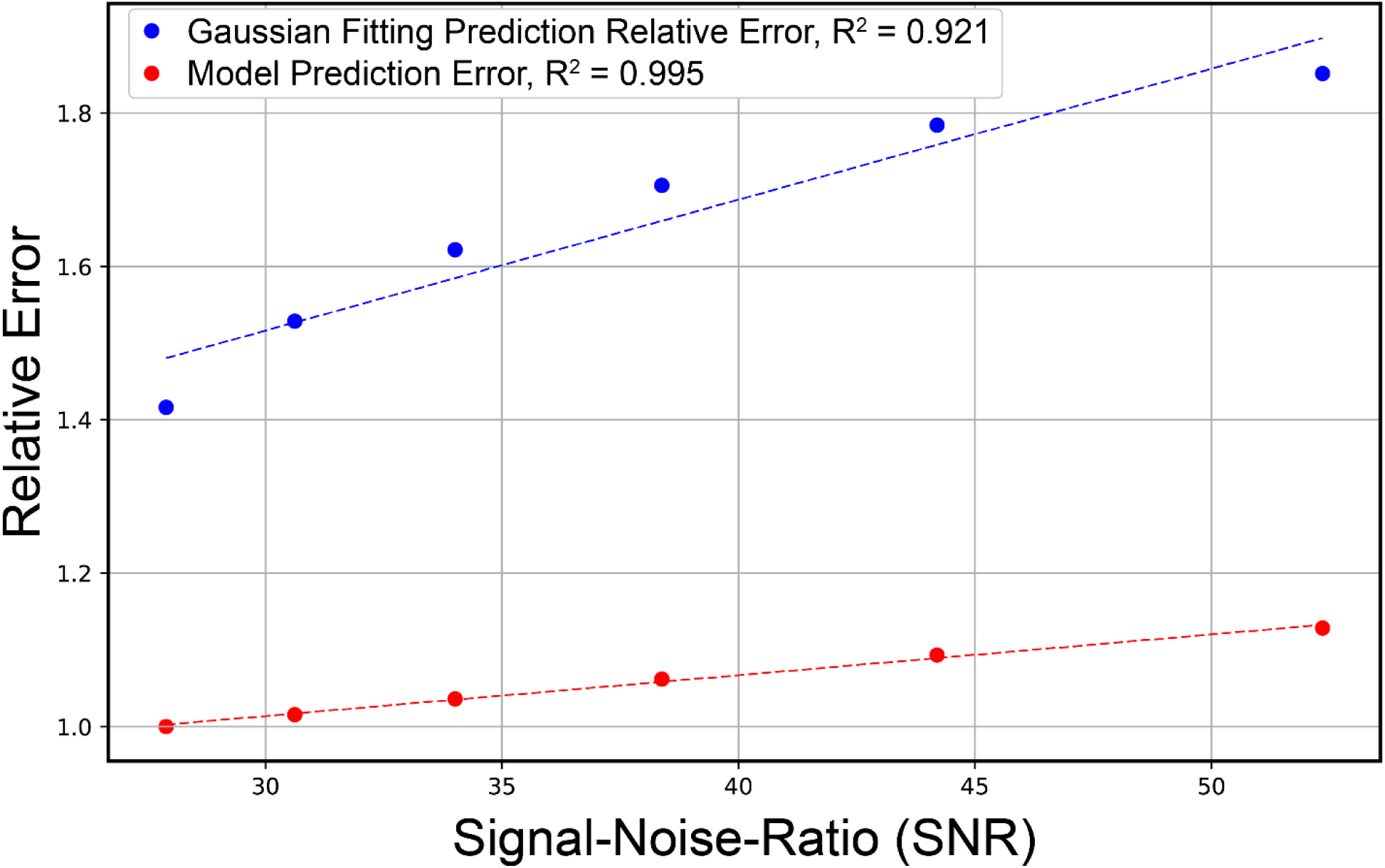
Comparison of trends for Relative MAE in AUC prediction vs signal-noise-ratio for both Gaussian fitting and the model.

Herein, we demonstrate a computational pipeline comprised of two separate neural network models for recovering the hyperspectral information content of IR imaging from sparsely sampled DFIR data. Our networks were trained using a combination of experimental and simulated data to capture the distribution of line shapes, widths, and noise found in spectra from a wide range of biological samples. While the up-scaling of spectral data on its own may not be a new concept, integrating this technique with a machine learning-based deconvolution method is an integral step forward in the analysis of IR imaging data. By requiring the measurement of only 7 key wavenumbers for the reconstruction of spectra, our first model can circumvent the data acquisition overhead that so often burdens traditional spectroscopic imaging techniques, allowing for the rapid investigation of even the largest specimens. Most efforts aimed at optimizing the experimental acquisition time in spectroscopic imaging have focused on coarser spatial sampling^63^, wherein only a subset of the pixels need to be sampled for reconstruction of the image at a given IR band. AI based tools have been employed to that end as well^63^. On the other hand, it is common practice to use some form of spectral deconvolution for hyperspectral data, such as those acquired in FTIR and spontaneous Raman imaging^64–66^. However, to the best of our knowledge, no AI based approach has been implemented towards coarsely sampling the spectral domain, as has been implemented in our work. While recording intensities at desired frequencies can suffice for well separated transitions in vibrational spectra, for bands that are convolutions of multiple transitions, such as protein vibrations this analysis becomes much more complex^48, 67^. The analysis of hyperspectral data in such cases requires additional deconvolution, typically through band fitting, and the resulting AUCs of the fitted components can be interpreted as proportional to corresponding concentrations or abundance. The AUC prediction model outlined here, presents clear advantages in data analysis time, prediction accuracy, and noise tolerance over traditional Gaussian fitting-based approaches to spectral deconvolution. Our model, despite using more sparsely sampled data than traditional Gaussian fitting, can evaluate spectral features like AUC values with higher accuracy. Because Gaussian fitting’s increased sensitivity to noise, our model also shows a better ability to adjust to lower SNR within spectral data. We note that the two models can be combined into a single model that simply predicts AUCs from seven spectral bands. However, we trained two separate models, because even individually they can benefit data acquisition and analysis approaches of different experimental implementations of IR imaging. For example, FTIR imaging systems, where the entire mid-IR spectra is acquired by design, can just utilize the second model for deconvolution. On the other hand, imaging systems capable of acquiring discrete frequencies (DFIR) can make use of only the first model for spectral reconstruction, or both models in conjunction when deconvolution is desired. The framework reported herein is useful for different modalities of IR imaging, and particularly for DFIR, where the time required data acquisition and analysis can be significantly reduced.

A key aspect of our approach is the potential for application towards other spectroscopic modalities and spectral regions. We have focused on the Amide I region (1600-1700 cm^-1^) due to its significant relevance in identifying protein secondary structures, making it immensely informative for studies related to protein conformation and interactions. We anticipate expansion of this framework beyond Amide vibrations to other spectral regions that are rich in molecular information, such as the 3-micron region (approximately 2500-3700 cm^-1^). C-H, O-H and N-H stretching modes from different molecular moieties such as lipids, proteins, nucleic acids etc. absorb in this spectral region. These bands are also convolutions of different molecular components, and our method can be easily applied to deciphering molecular compositions in these spectral regions as well. Secondly, our methodology is not limited to conventional IR spectroscopic imaging but is also generalizable to various forms of discrete frequency imaging techniques, such as photothermal IR imaging and Stimulated Raman Scattering (SRS) microscopy. These techniques are particularly beneficial in biological and medical research for their label-free and non-destructive nature, allowing for in vivo and dynamic studies with high specificity and sensitivity. It should be noted that successfully applying our approach to these modalities requires meticulous construction of the training dataset. It is essential that the dataset accurately represents the signal-to-noise ratio (SNR) inherent in the respective methods to ensure robust and reliable predictive performance. However, given our model leverages the fundamental variations in the Amide I spectra for spectral reconstruction and deconvolution, this approach should be directly extendable to imaging modalities that use mid IR spectra for chemical contrast, such as photothermal IR imaging. This also allows for wide applicability of this approach. Since our model is trained to recognize only the variations only in the spectral domain by using largely simulated spectra, it is not limited in the same way as networks trained on specific imaging systems would be in terms of transferability, which remains a key challenge in developing ‘turn-key’ solutions for data analysis in chemical imaging using machine learning. Our model is readily usable on all forms of IR data encompassing the Amide I region and can be easily retrained to be compatible with other spectral regions or methods such as Raman microspectroscopy. We aim to apply this model to other modalities of spectral imaging in future work.

It is important to note that the weights and biases of our network have been trained using largely simulated data, guided by the known and observed secondary structure distributions within the Amide I spectra of tissue samples. Specifically, the upscaling model is trained using a combination of synthetic and experimental data, while the AUC prediction model is trained on entirely simulated data. The spectra for training were simulated to capture same distribution of line shapes, widths, and noise as our experimental data. The reason for the choice of simulated data is the following. Firstly, it allows for sampling for a broader range of training data for the model without the burden of experimental data acquisition. Fitting and theoretical simulation of protein amide vibrations have been extensively studied and used in IR spectroscopy, which allows us to adopt this approach without any loss of generality. Of course, this may not be viable for all kinds of spectroscopic imaging techniques, where an extensive database for protein spectra does not exist. Secondly, in the case of experimental data, determining the ground truth (AUC) values requires Gaussian fitting, a process we aim to avoid. Gaussian fitting is time-consuming and plagued by the issues mentioned earlier, such as sensitivity to noise. Given the assumption that the spectra can be represented as the sum of three Gaussians, it is more suitable to consider this as the actual ground truth. In essence, our goal is to deduce the fundamental Gaussian parameters of the Amide I spectra. Of course, this does not apply to our first model, which only reconstructs spectra at higher frequency resolution from discrete bands, and hence we have used both simulated and experimental data to train this network. An important aspect to consider here is that Gaussian fitting of hyperspectral image data essentially deconvolutes each spectrum individually, without leveraging any information from the adjacent pixels corresponding to spectral location. Neural network based approaches, like the one used here, can be further optimized by factoring in the relationship of localized spectra to those from adjoining pixels whose properties are inherently related. This can potentially further the fidelity and applicability of these approaches towards analysis of complex spectral data in a wide variety of specimens, that would be otherwise challenging to address with conventional deconvolution strategies.

### Conclusions

In summary, we have introduced a novel approach that harnesses the power of machine learning to address challenges associated with acquiring and analyzing discrete frequency IR imaging. We show that a simple two-step regressive neural network model can be trained to use using spectral intensities at discrete frequencies to accurately replicate results from Gaussian curve fitting. The first network uses only seven wavenumbers to upscale spectra to higher resolutions, thus achieving a sixfold reduction in data acquisition time. This upscaled spectral data is then utilized by a second network to predict AUCs of the spectral components, akin to Gaussian fitting, but 3000 times faster. As a result, our approach drastically reduces both the time required for data acquisition and analysis, enabling the prediction of important variations in protein structure, which could be crucial for expanding the applications of discrete frequency imaging to proteinopathies. Since the networks are trained using largely simulated spectral data, they can be readily extended towards analyzing discrete frequency IR images from a wide variety of tissue specimens, including but not limited to brain sections from various neuro degenerative diseases. We note that our model reproduces both the amyloid positive and negative regions with high fidelity, which underscores the robustness of our approach. It should be noted that this study does not aim to classify spectral data as diseased or benign. Instead, it focuses on developing neural networks capable of reconstructing any amide I spectrum, regardless of its origin and relevance to disease, from a discrete set of bands and generating AUCs of constituent bands using a comprehensive library of simulated spectra. Thus, these models can have a broader application towards characterization of other diseases beyond Alzheimer’s, as studied here. Furthermore, our approach is independent of the spatial image parameters that may vary from different imaging systems and recognizes only the intensity variations arising from fundamental changes in protein secondary structure distributions. As a result, this also has the potential to be adapted for application for other spectroscopic modalities and spectral regions of interest, such as Raman spectroscopy. By tailoring the training datasets and expanding the application scope, our approach can substantially enhance the utility of spectroscopic imaging across various scientific fields, providing a powerful tool for detailed molecular characterization in diverse applications ranging from materials science to biomedical diagnostics.

## References

(1) Bhargava, R. Digital histopathology by infrared spectroscopic imaging. Annual Review of Analytical Chemistry 2023, 16 (1), 205–230.

(2) Pilling, M. J.; Henderson, A.; Gardner, P. Quantum cascade laser spectral histopathology: breast cancer diagnostics using high throughput chemical imaging. Analytical chemistry 2017, 89 (14), 7348–7355.

(3) Mukherjee, S. S.; Bhargava, R. Phasor Representation Approach for Rapid Exploratory Analysis of Large Infrared Spectroscopic Imaging Data Sets. Analytical Chemistry 2023, 95 (30), 11365–11374.

(4) Baker, M. J.; Trevisan, J.; Bassan, P.; Bhargava, R.; Butler, H. J.; Dorling, K. M.; Fielden, P. R.; Fogarty, S. W.; Fullwood, N. J.; Heys, K. A. Using Fourier transform IR spectroscopy to analyze biological materials. Nature protocols 2014, 9 (8), 1771–1791.

(5) Perl, D. P. Neuropathology of Alzheimer’s disease. Mt Sinai J Med 2010, 77 (1), 32–42. DOI: 10.1002/msj.20157.

(6) DeTure, M. A.; Dickson, D. W. The neuropathological diagnosis of Alzheimer’s disease. Mol. Neurodegener. 2019, 14 (32). DOI: 10.1186/s13024-019-0333-5.

(7) Confer, M. P.; Holcombe, B. M.; Foes, A. G.; Holmquist, J. M.; Walker, S. C.; Deb, S.; Ghosh, A. Label-Free Infrared Spectroscopic Imaging Reveals Heterogeneity of β-Sheet Aggregates in Alzheimer’s Disease. J. Phys. Chem. Lett. 2021, 12 (39), 9662–9671. DOI: 10.1021/acs.jpclett.1c02306.

(8) McNaught, K. S. P.; Olanow, C. W. Protein aggregation in the pathogenesis of familial and sporadic Parkinson’s disease. Neurobiology of aging 2006, 27 (4), 530–545.

(9) Hashimoto, M.; Rockenstein, E.; Crews, L.; Masliah, E. Role of protein aggregation in mitochondrial dysfunction and neurodegeneration in Alzheimer’s and Parkinson’s diseases. Neuromolecular medicine 2003, 4, 21–35.

(10) Wanker, E. E. Protein aggregation and pathogenesis of Huntingtons disease: Mechanisms and correlations. 2000.

(11) Hoffner, G.; Djian, P. Protein aggregation in Huntington’s disease. Biochimie 2002, 84 (4), 273–278.

(12) Rokad, D.; Ghaisas, S.; Harischandra, D. S.; Jin, H.; Anantharam, V.; Kanthasamy, A.; Kanthasamy, A. G. Role of neurotoxicants and traumatic brain injury in α-synuclein protein misfolding and aggregation. Brain research bulletin 2017, 133, 60–70.

(13) Cruz-Haces, M.; Tang, J.; Acosta, G.; Fernandez, J.; Shi, R. Pathological correlations between traumatic brain injury and chronic neurodegenerative diseases. Translational neurodegeneration 2017, 6, 1–10.

(14) Metzger, F. G.; Schopp, B.; Haeussinger, F. B.; Dehnen, K.; Synofzik, M.; Fallgatter, A. J.; Ehlis, A.-C. Brain activation in frontotemporal and Alzheimer’s dementia: a functional near-infrared spectroscopy study. Alzheimer’s Research & Therapy 2016, 8, 1–12.

(15) Von Bergen, M.; Barghorn, S.; Li, L.; Marx, A.; Biernat, J.; Mandelkow, E.-M.; Mandelkow, E. Mutations of tau protein in frontotemporal dementia promote aggregation of paired helical filaments by enhancing local β-structure. Journal of Biological Chemistry 2001, 276 (51), 48165–48174.

(16) Prusiner, S. B. Prion diseases and the BSE crisis. Science 1997, 278 (5336), 245–251.

(17) Prusiner, S. B. Prions. Proceedings of the National Academy of Sciences 1998, 95 (23), 13363–13383.

(18) Koo, E. H.; Lansbury Jr, P. T.; Kelly, J. W. Amyloid diseases: abnormal protein aggregation in neurodegeneration. Proceedings of the National Academy of Sciences 1999, 96 (18), 9989–9990.

(19) Ross, C. A.; Poirier, M. A. Protein aggregation and neurodegenerative disease. Nature medicine 2004,10 (Suppl 7), S10–S17.

(20) Álvarez-Marimon, E.; Castillo-Michel, H.; Reyes-Herrera, J.; Seira, J.; Aso, E.; Carmona, M.; Ferrer, I.; Cladera, J.; Benseny-Cases, N. Synchrotron X-ray Fluorescence and FTIR Signatures for Amyloid Fibrillary and Nonfibrillary Plaques. ACS Chem. Neurosci. 2021, 12 (11), 1961–1971. DOI: 10.1021/acschemneuro.1c00048.

(21) Liao, C. R.; Rak, M.; Lund, J.; Unger, M.; Platt, E.; Albensi, B. C.; Hirschmugl, C. J.; Gough, K. M. Synchrotron FTIR reveals lipid around and within amyloid plaques in transgenic mice and Alzheimer’s disease brain. Analyst 2013, 138 (14), 3991–3997. DOI: 10.1039/C3AN00295K.

(22) Surowka, A. D.; Pilling, M.; Henderson, A.; Boutin, H.; Christie, L.; Szczerbowska-Boruchowska, M.; Gardner, P. FTIR imaging of the molecular burden around Aβ deposits in an early-stage 3-Tg-APP-PSP1-TAU mouse model of Alzheimer’s disease. Analyst 2017, 142 (1), 156–168. DOI: 10.1039/C6AN01797E.

(23) Röhr, D.; Boon, B. D. C.; Schuler, M.; Kremer, K.; Hoozemans, J. J. M.; Bouwman, F. H.; El-Mashtoly, S. F.; Nabers, A.; Großerueschkamp, F.; Rozemuller, A. J. M.;, et al. Label-free vibrational imaging of different Aβ plaque types in Alzheimer’s disease reveals sequential events in plaque development. Acta Neuropathologica Communications 2020, 8 (1), 222. DOI: 10.1186/s40478-020-01091-5.

(24) Holcombe, B.; Foes, A.; Banerjee, S.; Yeh, K.; Wang, S.-H. J.; Bhargava, R.; Ghosh, A. Intermediate Antiparallel β Structure in Amyloid β Plaques Revealed by Infrared Spectroscopic Imaging. ACS Chemical Neuroscience 2023, 14 (20), 3794–3803.

(25) Ghimire, H.; Garlapati, C.; Janssen, E. A.; Krishnamurti, U.; Qin, G.; Aneja, R.; Perera, A. U. Protein conformational changes in breast cancer sera using infrared spectroscopic analysis. Cancers 2020, 12 (7), 1708.

(26) Li, N.; An, L.; Shen, J. Spectral fitting using basis set modified by measured B0 field distribution. NMR in Biomedicine 2015, 28 (12), 1707–1715.

(27) Andreeva, A.; Karamancheva, I.; Hendlich, M. Secondary Structural Analysis of Chloramphenicol Acetyltransferase Type I Using FTIR Spectroscopy. BULGARIAN CHEMICAL COMMUNICATIONS 2001, 33 (1), 9–15.

(28) DeOliveira, D. B.; Trumble, W. R.; Sarkar, H. K.; Singh, B. R. Secondary structure estimation of proteins using the amide III region of Fourier transform infrared spectroscopy: application to analyze calcium-binding-induced structural changes in calsequestrin. Applied spectroscopy 1994, 48 (11), 1432–1441.

(29) Thumanu, K.; Tanthanuch, W.; Lorthongpanich, C.; Heraud, P.; Parnpai, R. FTIR microspectroscopic imaging as a new tool to distinguish chemical composition of mouse blastocyst. Journal of Molecular Structure 2009, 933 (1-3), 104–111.

(30) Ishigaki, M.; Yasui, Y.; Puangchit, P.; Kawasaki, S.; Ozaki, Y. In vivo monitoring of the growth of fertilized eggs of medaka fish (Oryzias latipes) by near-infrared spectroscopy and near-Infrared imaging—a marked change in the relative content of weakly hydrogen-bonded water in egg yolk just before hatching. Molecules 2016, 21 (8), 1003.

(31) Siriwaseree, J.; Sanachai, K.; Aiebchun, T.; Tabtimmai, L.; Kuaprasert, B.; Choowongkomon, K. Synchrotron Fourier transform infrared microscopy spectra in cellular effects of Janus Kinase inhibitors on myelofibrosis cancer cells. ACS omega 2022, 7 (26), 22797–22803.

(32) Wang, D.; Ding, Y.-S.; Guo, Z.-H.; Min, S.-G. The application of near-infrared spectra micro-image in the imaging analysis of biology samples. Journal of Innovative Optical Health Sciences 2014, 7 (04), 1350062.

(33) Lam, F.; Liang, Z. P. A subspace approach to high-resolution spectroscopic imaging. Magnetic resonance in medicine 2014, 71 (4), 1349–1357.

(34) Blum, C.; Cesa, Y.; Escalante, M.; Subramaniam, V. Multimode microscopy: spectral and lifetime imaging. Journal of the Royal Society Interface 2009, 6 (suppl_1), S35-S43.

(35) He, H.; Yan, S.; Lyu, D.; Xu, M.; Ye, R.; Zheng, P.; Lu, X.; Wang, L.; Ren, B. Deep learning for biospectroscopy and biospectral imaging: state-of-the-art and perspectives. ACS Publications: 2021.

(36) Berisha, S.; Lotfollahi, M.; Jahanipour, J.; Gurcan, I.; Walsh, M.; Bhargava, R.; Van Nguyen, H.; Mayerich, D. Deep learning for FTIR histology: leveraging spatial and spectral features with convolutional neural networks. Analyst 2019, 144 (5), 1642–1653.

(37) Gurbani, S. S.; Schreibmann, E.; Maudsley, A. A.; Cordova, J. S.; Soher, B. J.; Poptani, H.; Verma, G.; Barker, P. B.; Shim, H.; Cooper, L. A. A convolutional neural network to filter artifacts in spectroscopic MRI. Magnetic resonance in medicine 2018, 80 (5), 1765–1775.

(38) Romeo, M. J.; Dukor, R. K.; Diem, M. Introduction to spectral imaging, and applications to diagnosis of lymph nodes. Vibrational Spectroscopy for Medical Diagnosis 2008, 1–26.

(39) Trajanovski, S.; Shan, C.; Weijtmans, P. J.; de Koning, S. G. B.; Ruers, T. J. Tongue tumor detection in hyperspectral images using deep learning semantic segmentation. IEEE transactions on biomedical engineering 2020, 68 (4), 1330–1340.

(40) Mei, S.; Jiang, R.; Li, X.; Du, Q. Spatial and spectral joint super-resolution using convolutional neural network. IEEE Transactions on Geoscience and Remote Sensing 2020, 58 (7), 4590–4603.

(41) Falahkheirkhah, K.; Yeh, K.; Mittal, S.; Pfister, L.; Bhargava, R. Deep learning-based protocols to enhance infrared imaging systems. Chemometrics and Intelligent Laboratory Systems 2021, 217, 104390.

(42) Aerts, H. J.; Velazquez, E. R.; Leijenaar, R. T.; Parmar, C.; Grossmann, P.; Carvalho, S.; Bussink, J.; Monshouwer, R.; Haibe-Kains, B.; Rietveld, D. Decoding tumour phenotype by noninvasive imaging using a quantitative radiomics approach. Nature communications 2014, 5 (1), 4006.

(43) Sutton, R. T.; Pincock, D.; Baumgart, D. C.; Sadowski, D. C.; Fedorak, R. N.; Kroeker, K. I. An overview of clinical decision support systems: benefits, risks, and strategies for success. NPJ digital medicine 2020, 3 (1), 17.

(44) Susi, H.; Byler, D. M. Protein structure by Fourier transform infrared spectroscopy: second derivative spectra. Biochemical and biophysical research communications 1983, 115 (1), 391–397.

(45) Rieppo, L.; Saarakkala, S.; Närhi, T.; Helminen, H.; Jurvelin, J.; Rieppo, J. Application of second derivative spectroscopy for increasing molecular specificity of fourier transform infrared spectroscopic imaging of articular cartilage. Osteoarthritis and cartilage 2012, 20 (5), 451–459.

(46) Confer, M. P.; Holcombe, B. M.; Foes, A. G.; Holmquist, J. M.; Walker, S. C.; Deb, S.; Ghosh, A. Label-free infrared spectroscopic imaging reveals heterogeneity of β-sheet aggregates in Alzheimer’s disease. The journal of physical chemistry letters 2021, 12 (39), 9662–9671.

(47) Tooke, P. Fourier self-deconvolution in IR spectroscopy. TrAC Trends in Analytical Chemistry 1988, 7 (4), 130–136.

(48) Yang, W.-J.; Griffiths, P. R.; Byler, D. M.; Susi, H. Protein conformation by infrared spectroscopy: resolution enhancement by Fourier self-deconvolution. Applied spectroscopy 1985, 39 (2), 282–287.

(49) Arrondo, J. L. R.; Iloro, I.; Garcia-Pacios, M.; Goñi, F. M. Two-dimensional infrared correlation spectroscopy. Advanced Techniques in Biophysics 2006, 73–88.

(50) Fabian, H.; Szendrei, G.; Mantsch, H.; Otvos, L. Comparative analysis of human and Dutch-type Alzheimer β-amyloid peptides by infrared spectroscopy and circular dichroism. Biochemical and biophysical research communications 1993, 191 (1), 232–239.

(51) Banerjee, S.; Holcombe, B.; Ringold, S.; Foes, A.; Ghosh, A. Structural heterogeneity of amyloid aggregates identified by spatially resolved nanoscale infrared spectroscopy. bioRxiv 2022, 2022.2005. 2007.491036.

(52) Mangialasche, F.; Kivipelto, M.; Solomon, A.; Fratiglioni, L. Dementia prevention: current epidemiological evidence and future perspective. Alzheimer’s research & therapy 2012, 4, 1–8.

(53) Alturkistani, H. A.; Tashkandi, F. M.; Mohammedsaleh, Z. M. Histological stains: a literature review and case study. Global journal of health science 2016, 8 (3), 72.

(54) Yang, J.; Liu, Z.; Wang, C.; Yang, R.; Rathkey, J. K.; Pinkard, O. W.; Shi, W.; Chen, Y.; Dubyak, G. R.; Abbott, D. W. Mechanism of gasdermin D recognition by inflammatory caspases and their inhibition by a gasdermin D-derived peptide inhibitor. Proceedings of the National Academy of Sciences 2018, 115 (26), 6792–6797.

(55) Yeh, K.; Kenkel, S.; Liu, J.-N.; Bhargava, R. Fast infrared chemical imaging with a quantum cascade laser. Analytical chemistry 2015, 87 (1), 485–493.

(56) DeVetter, B. M.; Kenkel, S.; Mittal, S.; Bhargava, R.; Wrobel, T. P. Characterization of the structure of low-e substrates and consequences for IR transflection measurements. Vibrational Spectroscopy 2017, 91, 119–127.

(57) Hardy, J.; Selkoe, D. J. The amyloid hypothesis of Alzheimer’s disease: progress and problems on the road to therapeutics. science 2002, 297 (5580), 353–356.

(58) Braak, H.; Braak, E. Neuropathological stageing of Alzheimer-related changes. Acta neuropathologica 1991, 82 (4), 239–259.

(59) Jack Jr, C. R.; Bennett, D. A.; Blennow, K.; Carrillo, M. C.; Dunn, B.; Haeberlein, S. B.; Holtzman, D. M.; Jagust, W.; Jessen, F.; Karlawish, J. NIA-AA research framework: toward a biological definition of Alzheimer’s disease. Alzheimer’s & dementia 2018, 14 (4), 535–562.

(60) Fellows, A. P.; Casford, M. T.; Davies, P. B. Spectral analysis and deconvolution of the amide I band of proteins presenting with high-frequency noise and baseline shifts. Applied Spectroscopy 2020, 74 (5), 597–615.

(61) Wilkosz, N.; Czaja, M.; Seweryn, S.; Skirlińska-Nosek, K.; Szymonski, M.; Lipiec, E.; Sofińska, K. Molecular spectroscopic markers of abnormal protein aggregation. Molecules 2020, 25 (11), 2498.

(62) Laumer, J.; O’Leary, S. K. A root-mean-square-error analysis of two-peak Gaussian and Lorentzian fittings of thin-film carbon Raman spectral data. Journal of Applied Physics 2019, 126 (4).

(63) Reihanisaransari, R.; Gajjela, C. C.; Wu, X.; Ishrak, R.; Corvigno, S.; Zhong, Y.; Liui, J.; Sood, A. K.; Mayerich, D.; Berisha, S. Rapid hyperspectral photothermal mid-infrared spectroscopic imaging from sparse data for gynecologic cancer tissue subtyping. ArXiv 2024.

(64) Belbachir, K.; Noreen, R.; Gouspillou, G.; Petibois, C. Collagen types analysis and differentiation by FTIR spectroscopy. Analytical and bioanalytical chemistry 2009, 395, 829–837.

(65) Petibois, C.; Gouspillou, G.; Wehbe, K.; Delage, J.-P.; Déléris, G. Analysis of type I and IV collagens by FT-IR spectroscopy and imaging for a molecular investigation of skeletal muscle connective tissue. Analytical and bioanalytical chemistry 2006, 386, 1961–1966.

(66) Delaney, J. K.; Zeibel, J. G.; Thoury, M.; Littleton, R.; Palmer, M.; Morales, K. M.; de La Rie, E. R.; Hoenigswald, A. Visible and infrared imaging spectroscopy of Picasso’s Harlequin musician: mapping and identification of artist materials in situ. Applied spectroscopy 2010, 64 (6), 584–594.

(67) Byler, D. M.; Susi, H. Examination of the secondary structure of proteins by deconvolved FTIR spectra. Biopolymers: Original Research on Biomolecules 1986, 25 (3), 469–487.

