## Supporting Information for "A Machine Learning-based Approach for Quantification of Protein Secondary Structures from Discrete Frequency Infrared Images"

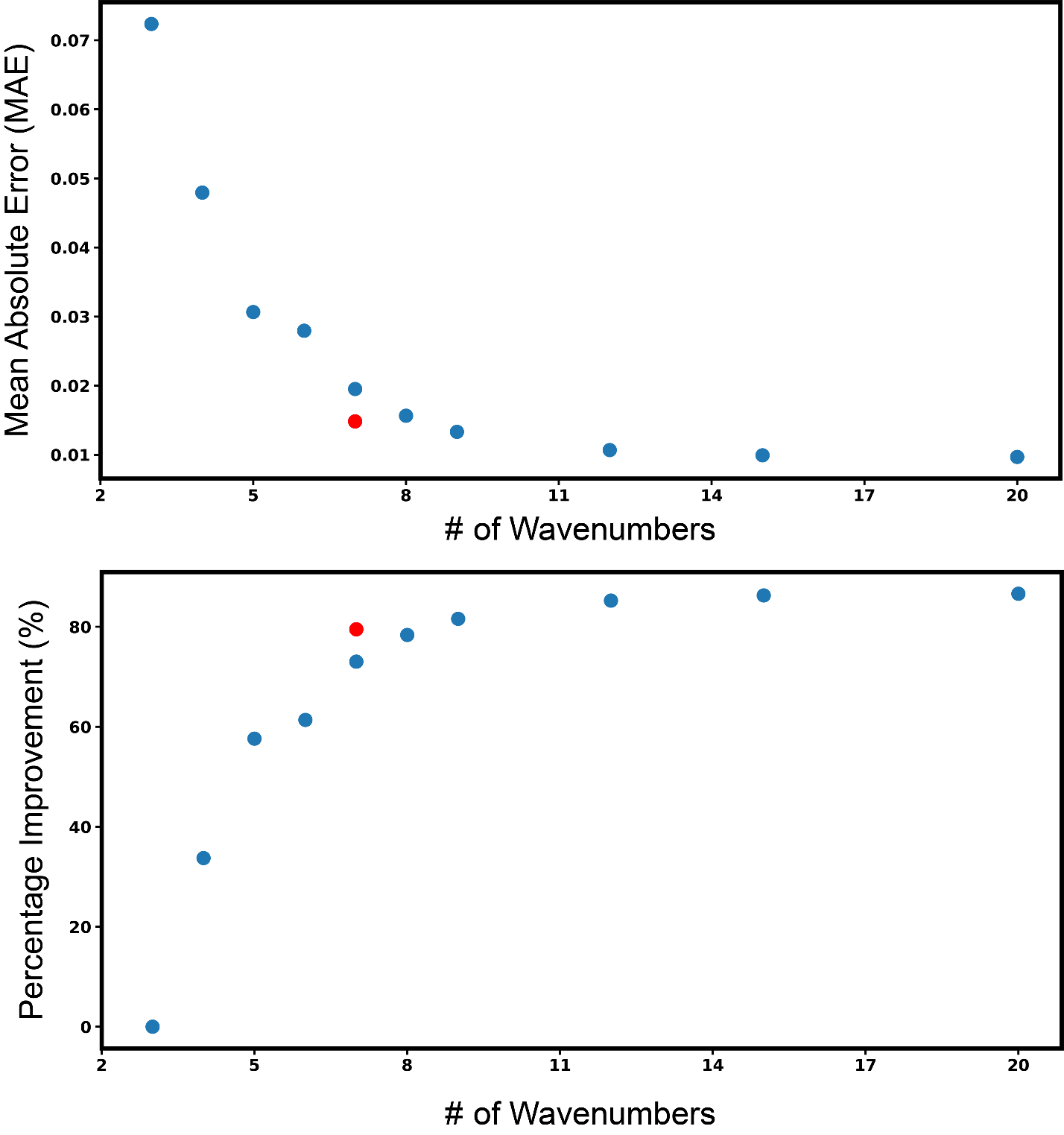

**Figure S1:** Relationship between the number of wavenumbers vs MAE (Top) and the relative percentage improvement in MAE (Bottom) of ANN1. The plot suggests a point of diminishing returns in model performance at around seven wavenumbers. The blue points represent wavenumber subsets that are equidistant from one another. The red point indicates a modified selection of seven wavenumbers for model training prioritizing significant spectral features.

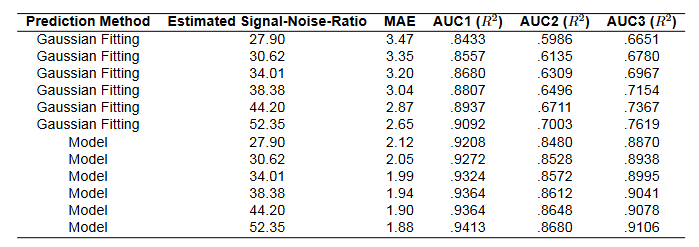

**Table S1:** Comparison of the model and Gaussian fittings performance when predicting AUC values with test spectra of varying random noise levels.

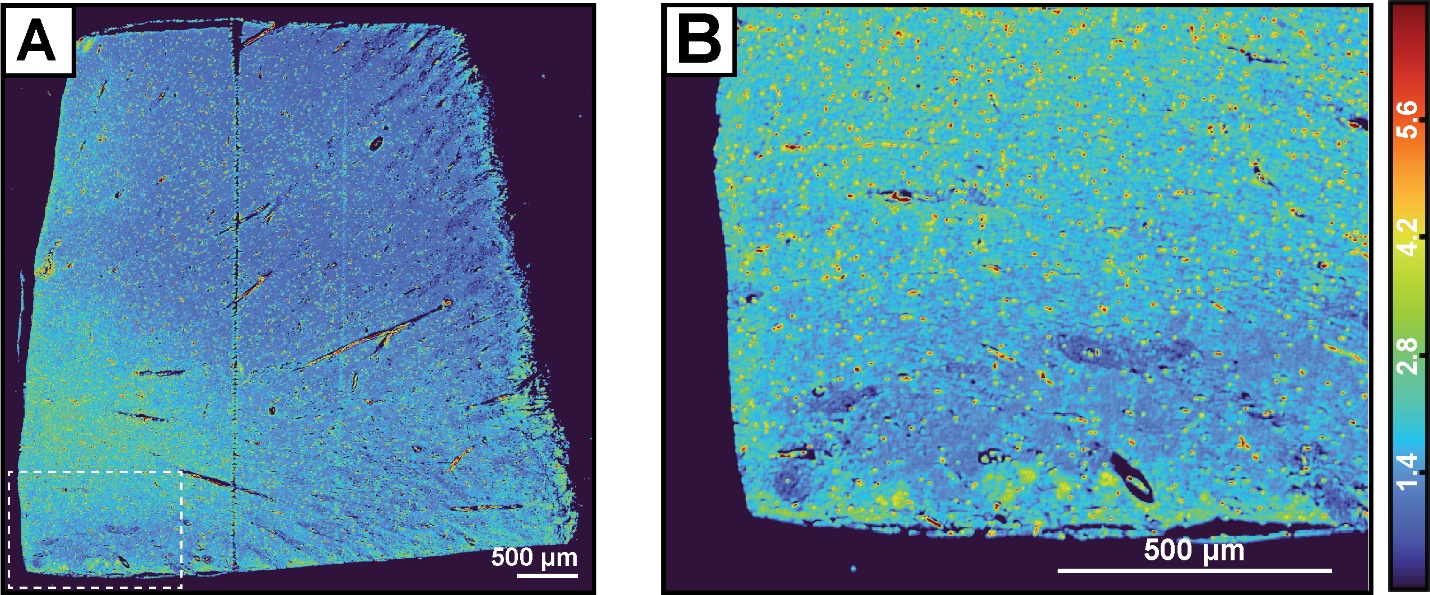

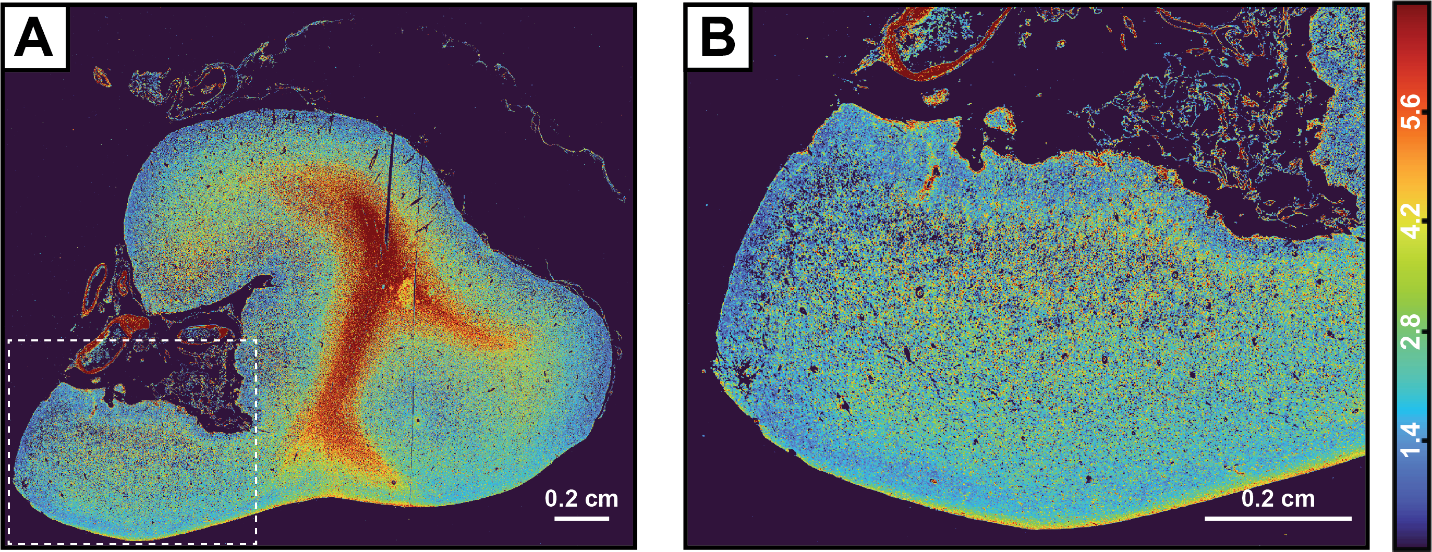
**Figure S2:** (A) Network predicted AUC1 image from collected hyperspectral data of a temporal lobe Alzheimer’s tissue section, (B) a smaller hypothetical region of interest denoted by a dashed white box.

**Figure S3:** (A) Network predicted AUC1 image from collected hyperspectral data of a frontal lobe Alzheimer’s tissue section, (B) a smaller hypothetical region of interest denoted by a dashed white box.

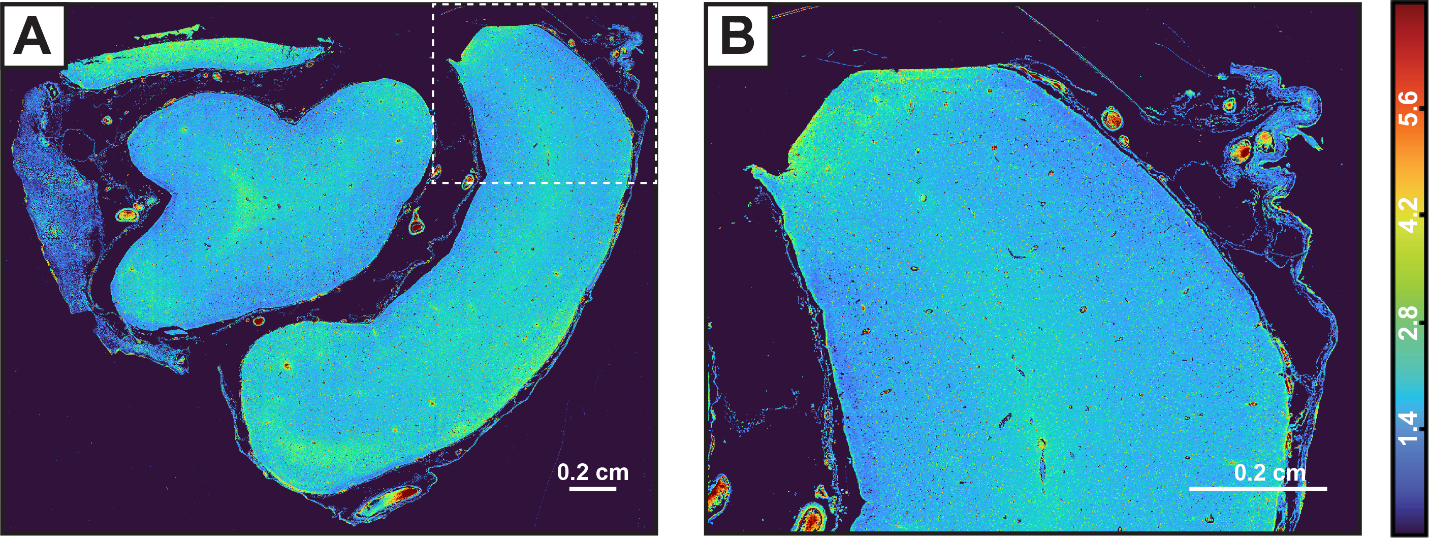

**Figure S4:** (A) Network predicted AUC1 image from collected hyperspectral data of a frontal lobe Alzheimer’s tissue section, (B) a smaller hypothetical region of interest denoted by a dashed white box.

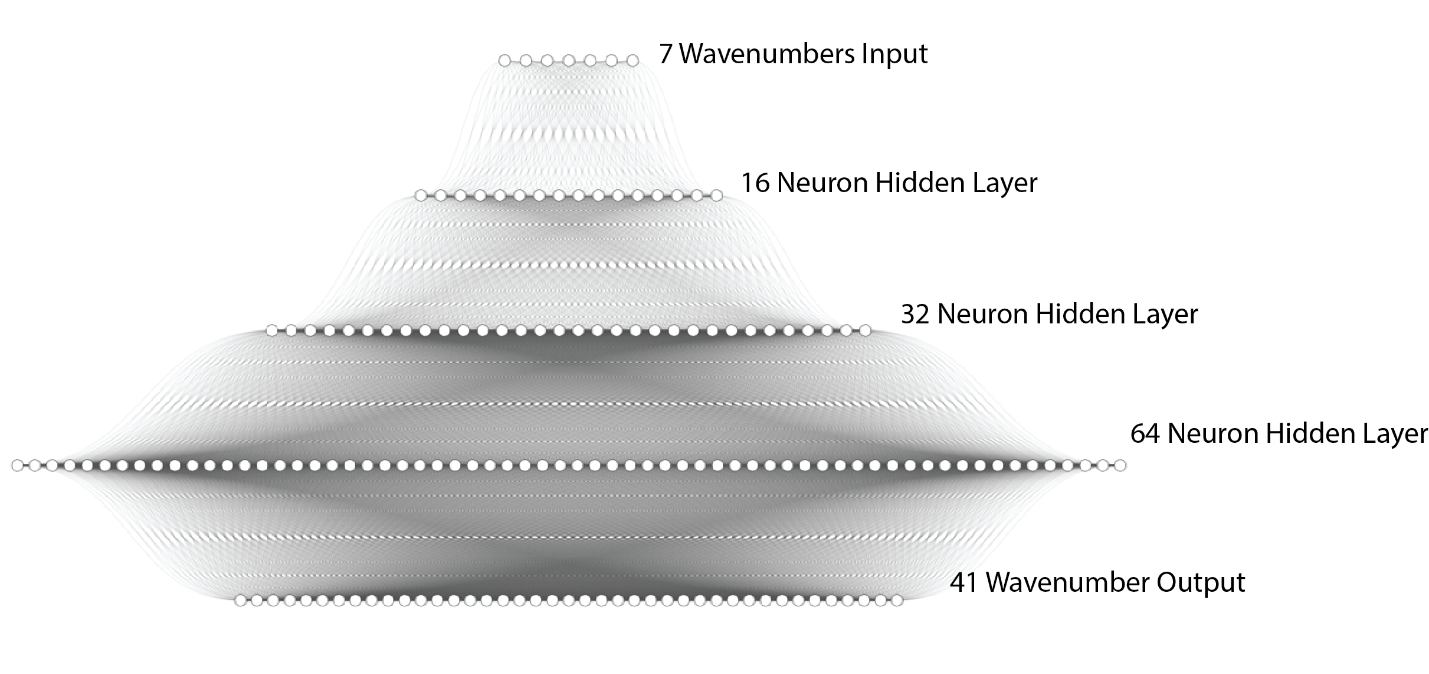

**Figure S5:** Spectral upscaling model (ANN1) architecture.

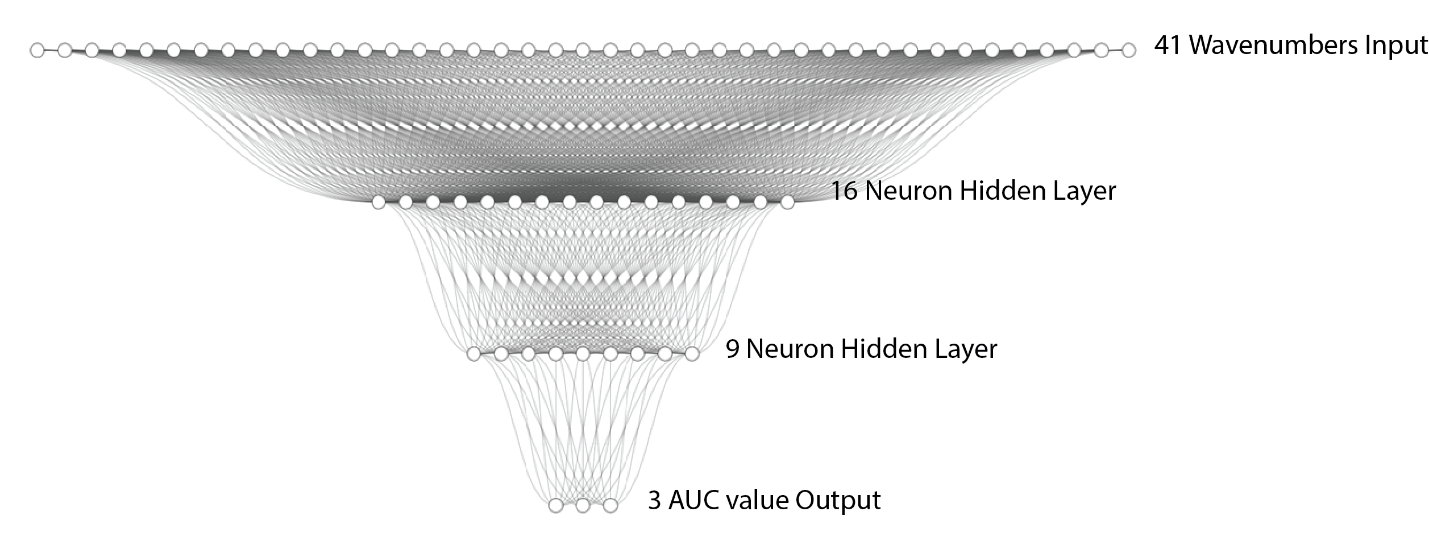

**Figure S6:** Spectral deconvolution model (ANN1) architecture.

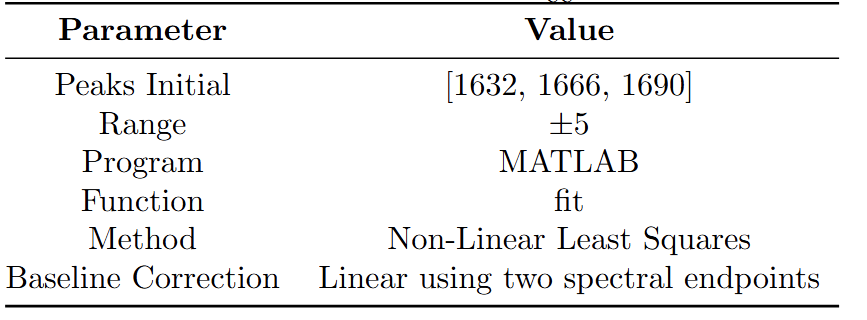

**Table S2:** Gaussian fitting parameters
